# Stress-responsive and metabolic gene regulation are altered in low S-adenosylmethionine

**DOI:** 10.1101/346205

**Authors:** Wei Ding, Daniel P Higgins, Dilip K. Yadav, Read Pukklia-Worley, Amy K Walker

## Abstract

S-adenosylmethionine (SAM) is the methyl donor that modifies proteins such as histones, nucleic acids and produces phosphatidylcholine. Thus variations in SAM levels could affect processes from lipogenesis to epigenetic gene regulation. SAM is hypothesized to link metabolism and chromatin modification, however, its role in acute gene regulation is poorly understood. We recently found that *Caenorhabditis elegans* with reduced SAM had deficiencies in bacterial-induced H3K4 trimethylation at selected pathogen-response genes, decreasing their expression and limiting survival on the pathogen *Pseudomonas aeruginosa*. This led us to the hypothesis that SAM may be generally required stress-responsive transcription. Here we show that *C. elegans* with low SAM fail to activate genome-wide transcriptional programs when exposed to bacterial or xenotoxic stress. However, heat shock responses were unaffected. We also investigated the role of two H3K4 methyltransferases that use SAM, *set-2/SET1*, and *set-16/MLL* and found that *set-2*/SET1 has a specific requirement in bacterial stress responses, whereas *set-16/MLL* was required for survival in all three stresses. These results define a role for SAM and H3K4 methyltransferases in the acute genome-wide remodeling of gene expression in response to stress. Finally, we find that the ability to modify metabolic gene expression correlates with enhanced survival in stress conditions.

## Introduction

Cellular functions are profoundly affected by metabolic state. For example, transcriptional regulation can be linked to metabolism through the modification of chromatin by methylation(Kaelin and McKnight 2013). Using the methyl groups produced by the 1-carbon cycle (1CC) and donated by S-adenosylmethionine (SAM), histone methyltransferases (HMTs) can change the regulatory state of the chromatin, promoting or limiting gene activity (Greer and Shi 2012). HMT activity can be controlled by recruitment of HMT containing complexes to specific genomic locations (Greer and Shi 2012). However, SAM availability may also affect histone methylation patterns (Walsh et al. 2018). SAM production by the 1-carbon cycle (1CC) and production can be affected by folate, methionine or choline levels or by other factors such as alcohol consumption (Mato et al. 2013). Variations in SAM levels have been proposed to mediate transgenerational inheritance of epigenetic patterns or other gene regulatory events (Mentch et al. 2015). However, a direct mechanistic connection has been difficult to establish (Mentch and Locasale 2016). Although SAM is necessary for all histone methylation events, *in vivo* studies have suggested that particular methylation marks are more sensitive to changes in SAM levels. For example, induced pluripotent stem cells, murine liver, and *C. elegans* all show decreases in H3K4me3 levels as SAM levels drop (Shyh-Chang et al. 2013; Kraus et al. 2014; Mentch and Locasale 2015). Furthermore, in budding yeast, a rise in SAM levels is followed by increases in H3K4me3 levels (Ye et al. 2017). Furthermore, these linked changes in SAM levels and H3K4 trimethylation also correlate with changes cell-type-specific gene expression and differentiation in iPS cells (Shyh-Chang et al. 2013). Finally, Dai et al. have recently demonstrated that treatments of high and low methionine, which is the precursor for SAM, alter H3K4me3 peak width at genes in steady state conditions in mouse liver and human cancer cells (Dai et al. 2018). Thus, SAM levels are tightly linked to H4K4me3 dynamics.

Trimethylation of H3K4 is a common modification occurring close to the start site of actively transcribed genes and is accomplished through the activity of the COMPASS complex (Shilatifard 2012). The KTM2 family of HMTs serves as the enzymatic activity of COMPASS providing mono, di and trimethylated states (Shilatifard 2006). In yeast, there is a single member, Set1, whereas mammals can use one of seven enzymes, within subfamilies of SET1, MLL (Mixed lineage leukemia) or THX (Trithorax) (Shilatifard 2012). However, the relationship between H3K4me3 and transcription is complex, as it does not appear to be necessary for gene expression in basal conditions (Lenstra et al. 2011). In yeast, Set1 has an important role in limiting the expression of ribosomal genes during the response to diamide stress (Weiner et al. 2012) suggesting that chromatin-modifying factors are especially critical when organisms experience stress. The H3K4 methyltransferase family in mammals appears to have overlapping as well as specialized functions in either specificity for mono, di or tri-methylation or through distinct roles in development (Shilatifard 2006). However, clearly defined roles for each MT have been difficult to discern.

*C. elegans* encoded a simplified KTM2 family containing three H3K4 methyltransferases, *set-2*/SET1, *set-16*/MLL and *ash-2*/THX (Wenzel et al. 2011). Interestingly, these methyltransferases have distinct developmental and tissue-specific biological functions. *set-2*/SET1 is broadly important for H3K4 trimethylation in embryos and the germline (Li and Kelly 2011; Xiao et al. 2011) and the intestine (Ding et al. 2015). Also, loss or reduction in *set-2*/SET1 transgenerationally influences sterility (Robert et al. 2014), lifespan (Greer et al. 2010) and lipid accumulation (Han et al. 2017). *ash-2* acts through the germline to affect lifespan and lipid accumulation in the intestine (Greer et al. 2010; Greer et al. 2011; Han et al. 2017). *set-16*/MLL, on the other hand, appears to be dispensable for H3K4me3 in the early embryo and germline (Li, 2011), while we found that it has a partial requirement in the adult intestine (Ding et al. 2015). Thus, while H3K4me3 marks the start sites of actively transcribed genes, the methyltransferases producing it can have diverse and long-acting biological effects.

Using a *C. elegans* model of low SAM, we previously found that transcriptional responses response to a bacterial pathogen failed and these bacterial-response genes did not show normal H3K4me3 close to the transcriptional start sites, (Ding et al. 2015). We also found the HMT *set-16*/MLL was required for full induction, whereas *set-2*/SET1 appeared dispensable (Ding et al. 2015). We hypothesized that animals with low SAM might fail to transcriptionally respond to stress and that the HMTs may also have distinct roles in modulating stress responses. In our present study, we set out to compare induction of transcriptional responses and survival upon stress exposure between *C. elegans* with reduced SAM (*sams-1(RNAi)*) and animals with limited H3K4me3 function, *set-2*/SET1, and *set-16*/MLL RNAi. Because distinct stresses may rely on different transcriptional activation mechanisms, we also compared whole-genome expression patterns in three stresses: pathogenic bacteria (PA14), xenotoxic (R24) and heat. We find that genes in the pathogen and xenotoxic stress response were indeed diminished in low SAM, with concomitant reductions in survival in these animals. However, while pathogen and xenotoxic-stress genes were affected both *set-2*/SET1 and *set-16*/MLL, *set-16*/MLL was uniquely required for survival in all three stresses. This suggests SAM and *set-16* have essential functions in transcriptional responses to these stresses. Interestingly, induction of heat stress genes, which are controlled primarily by promoter pausing of RNA Pol II (Adelman and Lis 2012), occurs even in low SAM and after H3K4 MT knockdown. Finally, we find that in addition to stress-responsive genes, regulation of metabolic genes may be key to the survival of animals with deficient H3K4 methylation during stress.

## RESULTS

### Large-scale changes in stress-induced gene expression in low SAM

Gene regulatory events can be controlled by histone methylation; however, it is not clear how levels of the methyl donor SAM may alter methylation patterns and gene expression in different physiological conditions (**Figure 1A**). In *C. elegans*, we previously found animals with a mutation in the SAM synthase *sams-1*, which have 50% of the SAM of wild-type animals (Walker et al. 2011), had poor survival upon the bacterial pathogen *Pseudomonas aeruginosa PA14 (Pseudomonas)* (Ding et al. 2015). SAM deficient animals failed to upregulate selected pathogen-response genes and had reduced global H3K4 trimethylation in intestinal nuclei as well at specific pathogen-response genes (Ding et al. 2015). We hypothesized this could represent a general failure of stress-responsive gene expression, as low SAM levels were unable to support rapid remodeling of H3K4 methylation as transcriptional needs changed. To test this model (**Figure 1B**), we used RNAi to knockdown *sams-1* or the H4K4me3 methyltransferases (HMTs) *set-2* or *set-16* that use SAM, then exposed animals to three stresses: bacterial (*Pseudomonas aeruginosa*, PA14), xenotoxic, or heat. For xenotoxic stress, we used R24, an agent that stimulates an immune and detoxification response in *C. elegans* (Pukkila-Worley and Ausubel 2012; Pukkila-Worley et al. 2014; Cheesman et al. 2016). Next, we used whole genome RNA sequencing to determine which genes changed in each stress and how they were affected by low SAM or depletion of the HMTs. Heat maps showing genes that greater than 2 fold in any of the conditions with a false discovery rate (FDR) of <0.01 showed distinct clusters for each stress (**Figure 1C**). For example, bacterial stress-induced genes grouped in three major clusters (a, b, c, e); a and c showed some overlap with heat stress-induced genes while b was largely non-overlapping with genes upregulated in controls from xenotoxic agent-induced or heat stress (**Figure 1C**). Interestingly, genes upregulated in PA14 cluster e appear reciprocally regulated during heat stress. Within bacterial or xenotoxic agent-induced stresses, *sams-1* appeared to have lower induction within the major clusters, whereas *set-2* and *set-16* appear to have intermediate effects (**Figure 1C**). Major clusters within heat shock-induced genes did not appear to share these large-scale changes. Thus, animals with low SAM appear to have distinct genome-wide expression patterns in response to pathogenic, xenotoxic or heat stress.

**Figure 1:**
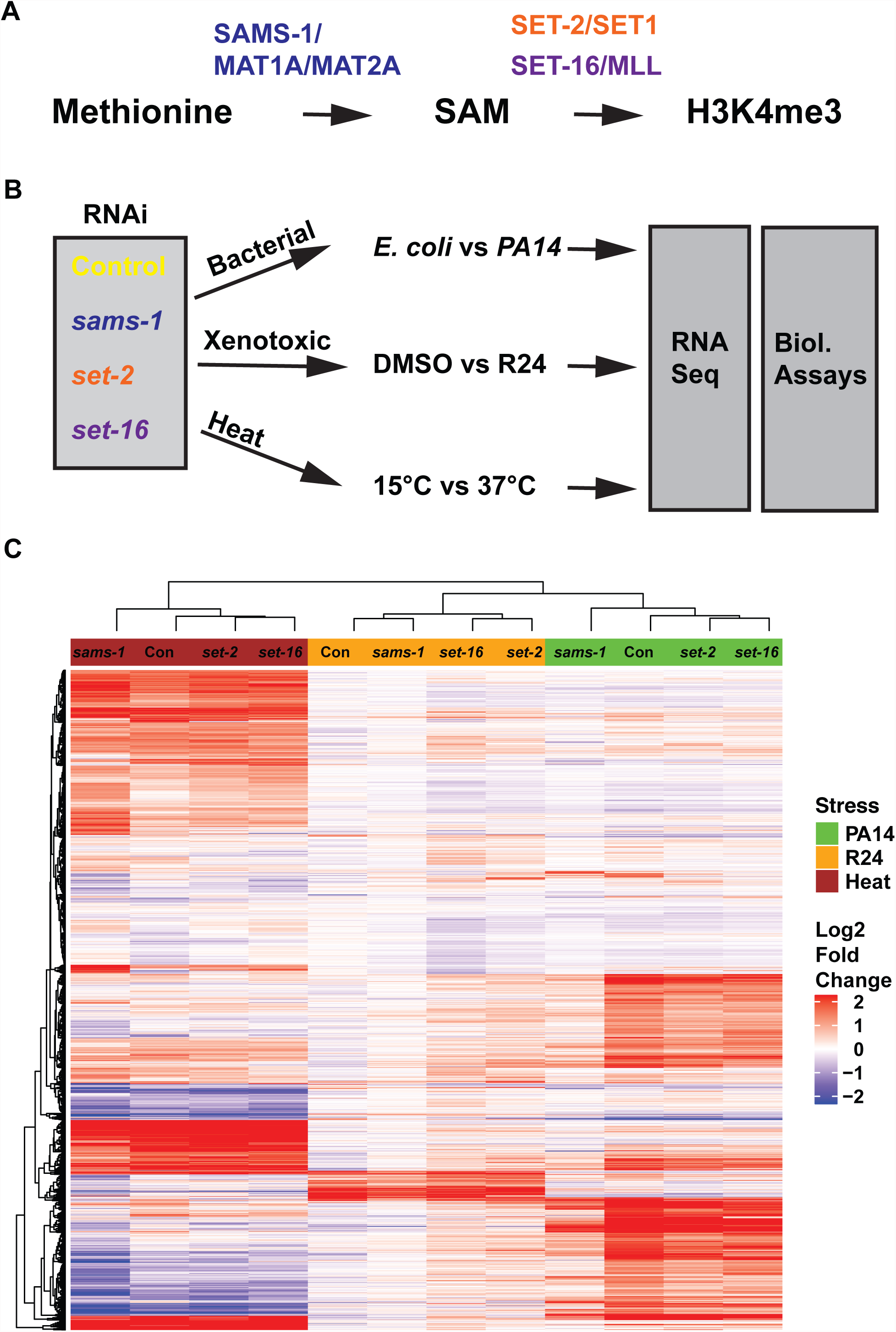
SAM plays an important role in the transcriptional response to stress. A. Schematic showing the metabolic link between s-adenosylmethionine (SAM) production and histone methylation **B.** Diagram of the experimental rationale comparing knockdowns of the SAM synthase *sams-1* and the H3K4 methyltransferases *set-2*/SET1 and *set-16*/MLL in the transcriptional response to three distinct stresses. **C.** Heat map showing unbiased hierarchical clustering of genes regulated 2 or more fold with an FDR of less than 0.01 in each of the stresses.

### Role of SAM in gene expression and H3K4 methylation in basal conditions

To understand the role of SAM in stress-induced gene expression in more detail, we first examined the role of SAM basal conditions, comparing genes induced between control (OP50) and pathogenic (*Pseudomonas aeruginosa* PA14) bacteria. In basal conditions, several hundred genes changed by more than two-fold after *sams-1(RNAi)* (**Supplemental_Figure_1B**), with significant overlap with our previous microarray results (Ding et al. 2015). We hypothesize that most of these gene expression changes were linked to methylation-dependent PC production, as they were returned to wild-type levels when PC levels were rescued by dietary choline (**Supplemental_Figure_1C**) (Ding et al. 2015). Both H3K4 methylation and PC production are major consumers of SAM (Walsh et al. 2018). Recently Ye, et al. show that H3K4 tri-methylation can increase in *Saccharomyces cerevisiae* when PC production is blocked and SAM levels increase (Ye et al. 2017). In agreement with these findings, we also observed that global H3K4me3 levels increase when the PC-producing methyltransferases *pmt-2* was knocked down (**Supplemental_Figure_1D**). Thus, in basal conditions, gene expression changes to compensate for decreases in PC are the predominant effect of *sams-1* loss, with negligible effects due to other methylation pathways. Finally, modified H3K4 may exist in several methylation states (Shilatifard 2006). Using immunostaining with antibodies to H3K4me1 and H3K4me2, we found that levels did not decease as they had with H3K4me3 (**Supplemental_Figure_1D**; (Ding et al. 2015)), suggesting that the trimethyalted state is most sensitive to SAM levels in adult *C. elegans* intestine.

### SAM is important for the transcriptional response to a bacterial stress

Next, we compared genes upregulated during the response to *Pseudomonas*. Control animals upregulated 651 genes more than two-fold in response to the bacterial stress (**Figure 2A; 2B, yellow; Supplementa_Table_1**) with a high concordance to previous studies identifying *Pseudomonas*-response genes (Troemel et al. 2006; Wong et al. 2007) (**Supplemental_Table_1**). Heat maps comparing genes upregulated more than 2-fold with an FDR of < 0.01 show lower induction after *sams-1* RNAi, with intermediate effects after *set-2* or *set-16* knockdown (**Figure 2A**). Focusing on the top 20 expressed genes in control animals, we find a significant reduction in expression (**Figure 2B**). Volcano plots comparing genome-wide expression patterns in control or *sams-1* RNAi after bacterial stress exposure also confirm a striking loss of the transcriptional response (**Figure 2C**) and finally, we find that few genes outside the pathogen response are induced after *sams-1* RNAi (**Figure 2D**). The transcriptional response to *Pseudomonas* also includes downregulation of a small subset of genes (Troemel et al. 2006). Comparisons between control and *sams-1(RNAi)* gene expression patterns show that a proportion of the two 20 downregulated genes in control animals fail to decrease after *sams-1* RNAi (**Supplemental_Figure_3A**) and that only about 5 percent of these genes overlap (**Supplemental_Figure_3B**). Thus, this whole genome data confirms our analysis of selected *Pseudomonas*-responsive genes (Ding et al. 2015) and shows that SAM is essential for the broad transcriptional changes occurring during stress caused by a pathogenic bacteria, reducing both total numbers of regulated genes and their magnitude.

**Figure 2:**
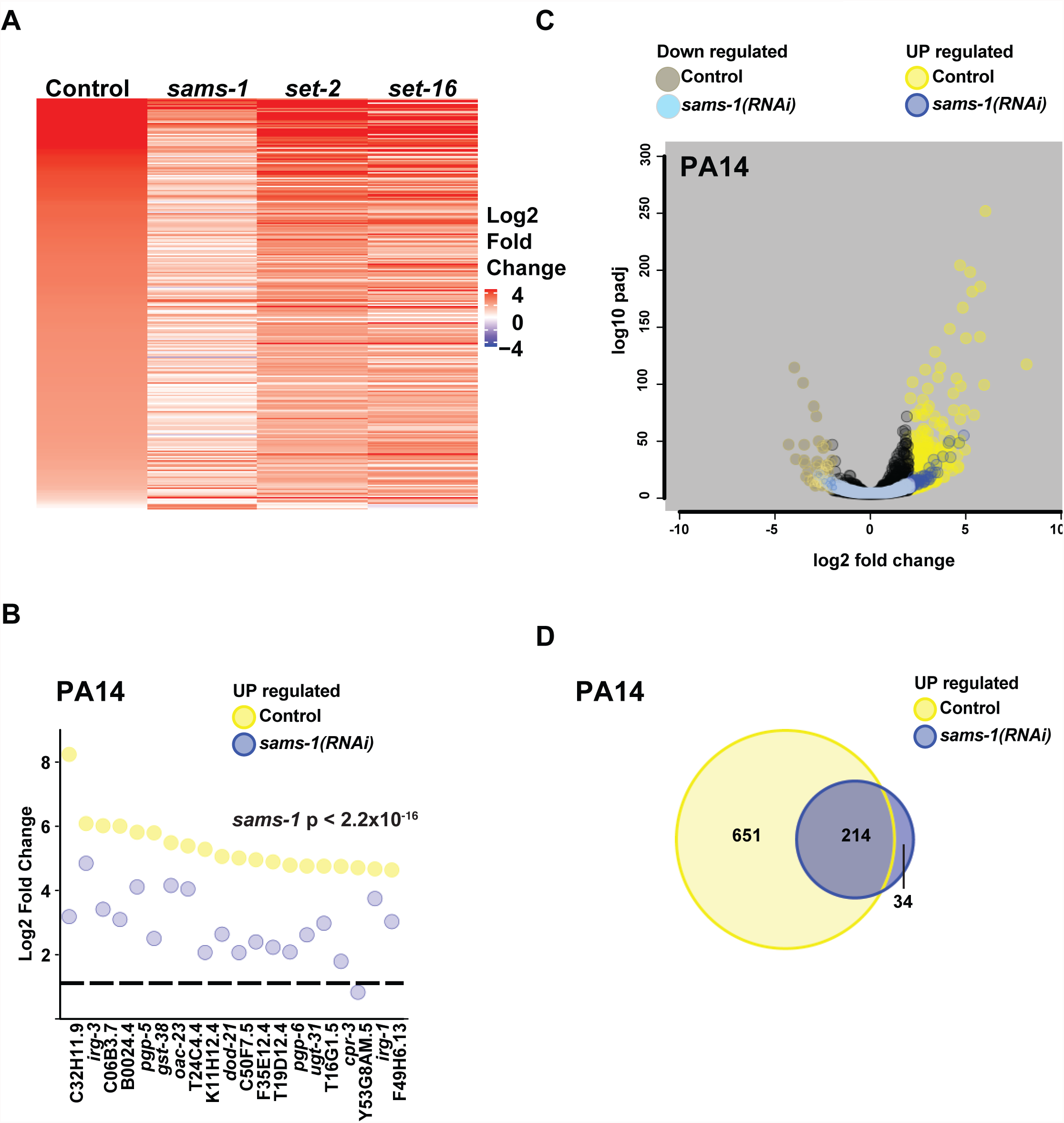
Transcriptional response to *Pseudomonas* requires *sams-1*. (**A**) Heat map showing genes upregulated by more than 2-fold with an FDR of less than 0.01 in *C. elegans* exposed to *Pseudomonas*. (**B**) Strip-plot comparing the top 20 genes upregulated in control vs. *sams-1(RNAi)* animals exposed to *Pseudomonas* (PA14). The dotted line is placed at one on the Y-axis. Statistical significance calculated by KS value. (**C**) Volcano plot of RNA sequencing data showing a reduced transcriptional response of *sams-1(RNAi)* animals to *Pseudomonas*, compared to controls. (**D**) Venn Diagram comparing overlap between genes upregulated more than 2-fold in control vs. *sams-1(RNAi)* animals exposed to *Pseudomonas*. Significant genes in RNA sequencing data (B-D) were from duplicate experiments and showed more than two fold changed with an FDR of <0.01. For B-D: Control: yellow, Up; brown: down; *sams-1(RNAi)* blue, Up; light blue, down.

### Transcriptional responses and survival upon xenobiotic challenge are diminished in low SAM

To determine how *sams-1* RNAi animals respond to a distinct stress, we treated control and *sams-1* RNAi animals with R24, a xenotoxic agent that induces both a cytochrome P450 and immune defenses dependent on a mitogen-activated kinase (MAPK) pathway in *C. elegans* (Pukkila-Worley and Ausubel 2012; Pukkila-Worley et al. 2014; Cheesman et al. 2016). Paradoxically, this gene activation program protects *C. elegans* exposed to *Pseudomonas* but is toxic to animals in basal laboratory conditions (Pukkila-Worley and Ausubel 2012; Cheesman et al. 2016). To identify and compare gene expression changes in low SAM animals exposed to this xenotoxic agent, we performed RNA-seq on control and *sams-1(RNAi)* after 6 hours of treatment with R24. We found that a robust set of genes was significantly upregulated by more than two-fold in control animals consistent to published results (Pukkila-Worley and Ausubel 2012; Pukkila-Worley et al. 2014; Cheesman et al. 2016), importantly, this set was substantially less regulated in *sams-1(RNAi)* animals (**Figure 3A**). The 20 most highly induced genes after R24 treatment included multiple cytochrome p450s as well as previously identified pathogen response genes (**Figure 3B, yellow**) (Pukkila-Worley and Ausubel 2012). Strip plots comparing levels in control and *sams-1(RNAi)* animals show that each gene was markedly decreased (**Figure 3C**). Volcano plots show that the genome-wide changes after treatment with the xenotoxic agent R24 largely failed to occur after *sams-1 RNAi* (**Figure 3C**). We also found that most of the genes induced in *sams-1* animals were part of the response to R24 in control animals (**Figure 3D**). R24 also induces the downregulation of a limited subset of genes (Pukkila-Worley and Ausubel 2012). Comparison between genes downregulated in control animal or after *sams-1* RNAi shows that *sams-1* RNAi also limits this downregulation (**Supplemental_Figure_3C, D**). Thus, low SAM attenuates the transcriptional response to a xenotoxic agent, just as it does to bacterial stress-responsive gene expression induced by *Pseudomonas*.

**Figure 3:**
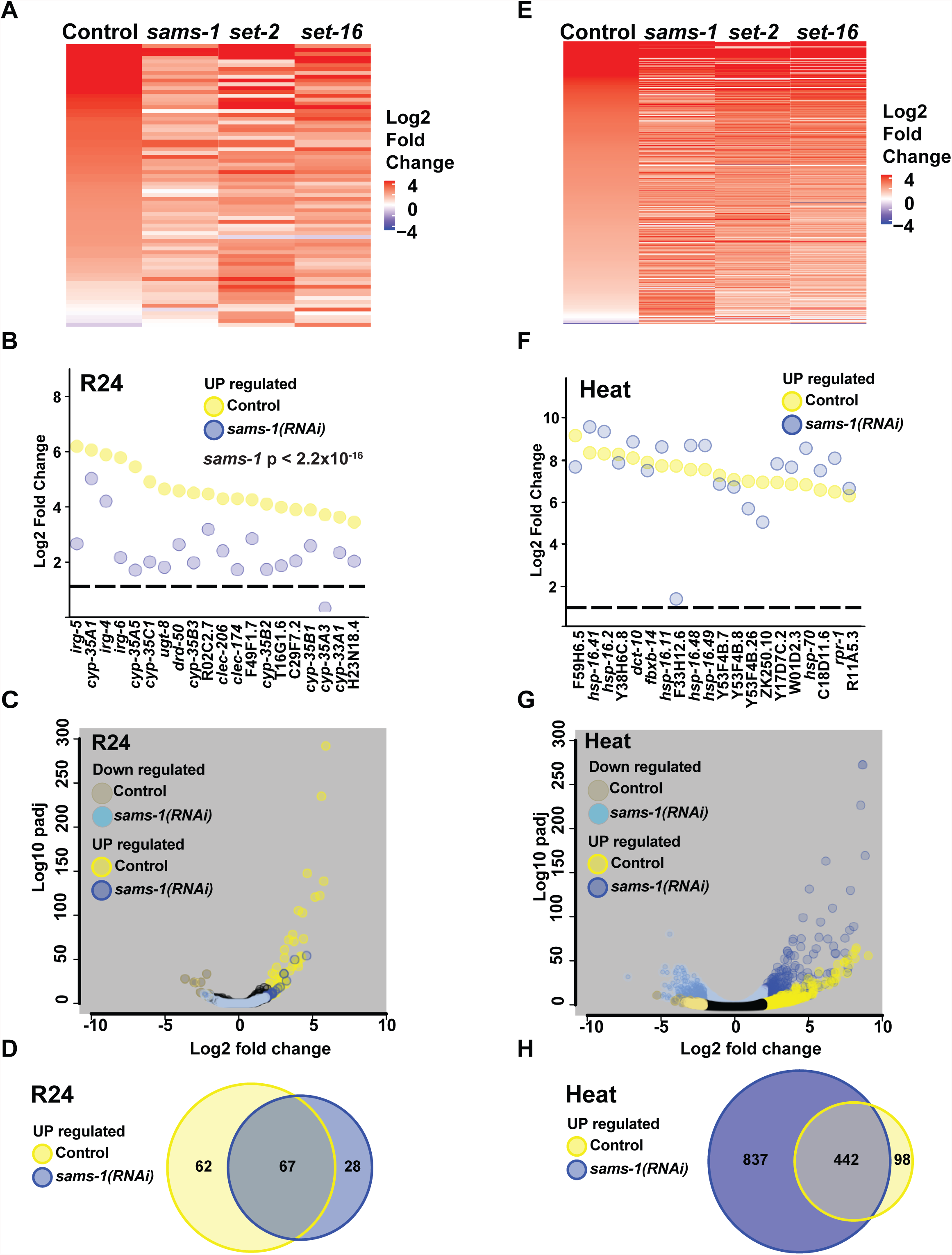
Differential transcriptional responses to a xenotoxic stress and heat stress after *sams-1(RNAi)* (**A**) Heat map showing genes upregulated by more than 2- fold with an FDR of less than 0.01 in *C. elegans* exposed to R24. **(B)** Strip plots showing that the top 20 genes upregulated in controls in response to R24 decreased are in *sams-1(RNAi)* animals. (**C**) Volcano plots of RNA sequencing data showing a reduced number of genes responding to the xenotoxic stress from R24 in *sams-1* animals vs. controls. (**D**) Venn diagrams show that the majority of genes upregulated more than two-fold in *sams-1* animals in response to R24 are also upregulated in controls. (**E**) Heat map showing genes upregulated by more than 2-fold with an FDR of less than 0.01 in *C. elegans* exposed to heat. (**F**) Strip-plot shows that the top 20 genes expressed after heat shock are upregulated similarly in control and *sams-1(RNAi)* animals. (**G**) Volcano plots show many genes outside of the heat shock response are changed in *sams-1(RNAi)* animals compared to controls. (**H**) Venn Diagram comparing gene sets regulated after heat stress shows many ectopic genes upregulated in *sams-1(RNAi)* animals. For A-H: Control: yellow, up; brown: down; *sams-1(RNAi)* blue, up; light blue, down. Significant genes in RNA sequencing data (B-H) were from duplicate experiments and showed more than two-fold changed with an FDR of <0.01.

### Heat shock transcriptional response occurs independently of SAM

Transcriptional response to bacteria and xenotoxic agents are predicted to follow a classic signal transduction pathway where the extracellular stimulus activates a cellular signaling pathway linked to individual transcription factors and upregulation of stress-specific gene expression (Estruch 2000). However, other stress-responsive genes expression, such as the heat shock genes, are regulated differently. RNA Pol II is paused at promoters of many heat shock genes and released into its elongating form in response to heat (Vihervaara et al. 2018). To determine if low SAM had the same effects on heat shock-dependent transcription as the bacterial or xenotoxic stress, we performed whole genome RNA sequencing on control, *sams-1*, *set-2* and *set-16* RNAi animals exposed to 37°C for one hour. We found that heat shock genes were strongly induced in control animals in these conditions (**Figure 3E-H**). In contrast to the bacterial or xenotoxic stress responses, comparison of control and *sams-1* patterns for genes induced at least 2-fold shows similar patterns(**Figure 3E**), suggesting that reduced SAM availability does not compromise the activation of heat-stress induced genes. Strip plots comparing expression of the top 20 genes activated in control animals compared to *sams-1* shows that most of the highly expressed genes are similarly or more highly expressed after *sams-1* RNAi (**Figure 3F**). In addition, volcano plots showing genome-wide comparisons of genes induced by heat shock in control or *sams-1* animals show a larger transcriptional response in *sams-1* animals, suggesting de-repression or activation of genes outside the canonical response (**Figure 3G**). Finally, as in the volcano plots, Venn diagrams confirm that the genes upregulated after heat shock in controls are also upregulated in *sams-1* animals and that many genes ectopic to the heat shock response also increase (**Figure 3H**). Control animals downregulated approximately 300 genes after heat shock; strikingly, nearly 2000 genes decreased in parallel *sams-1* RNAi animals (**Supplemental_Figure_3E, F**). Thus, while the canonical heat shock response seems to occur independently of SAM, other genes dramatically change during the transcriptional response to heat in low SAM.

### Differential attenuation of stress-responses after knockdown of the H3K4 methyltransferases *set-2*

Histone methyltransferases use SAM to modify specific histone residues, modifying the chromatin environment to provide distinct gene regulatory states. Yeast contain a single H3K4 HMT, which provides mono, di and trimethylated states (Shilatifard 2012) and functions within the COMPAS HMT complex. Mammals encode 7 H4K4 HMTs that have different specificity for methylation states (Shilatifard 2012). However, the non-redundant biological functions have been difficult to discern. *C. elegans* contains 3 H3K4 MTs that affect H4K4me3, *set-2*/SET1, *set-16*/MLL and *ash-2*/THX. These HMTs affect embryonic and germline development (Li and Kelly 2011; Xiao et al. 2011) and transgenerational inheritance through the germline (Greer et al. 2010; Greer et al. 2011). In our previous studies, we investigated the roles of *set-2*/SET1 and *set-16*/MLL in the adult *C. elegans* intestine, which is a key tissue in the pathogen response (Millet and Ewbank 2004). Because H3K4me3 has been associated with dynamically transcribed genes and our previous results showing increases in H3K4 tri-methylation at promoters of selected *Pseudomonas* responsive genes during infection (Ding et al. 2015), we sought to determine if *set-2*/SET1 or *set-16*/MLL were downstream of SAM-dependent responses during the stress response. In parallel with *sams-1(RNAi)* RNA experiments (**Figure 1-2**), we exposed *set-2(RNAi)* animals to *Pseudomonas*, the xenotoxic agent R24 or heat stress, extracted RNA and performed RNA-sequencing. Unbiased hierarchical clustering analysis of all genes significantly upregulated by any of these stresses, showed that *set-2* and *set-16* RNAi grouped within each stress, suggesting similar overall gene expression patterns (**Figure 1C**). Next, we used the same computational tools as in the *sams-1* analysis to compare expression patterns of *Pseudomonas* response genes after *set-2* RNAi. Interestingly, although heat maps show that activation bacterial-stress responsive genes are clearly diminished after *set-2* RNAi, the effect is less severe than in *sams-1(RNAi)* animals (**Figures 2A; 4A-D**). Direct comparison of the 20 most highly expressed genes shows reduced expression of several genes in *set-2* animals, in line with an intermediate effect between controls and *sams-1* (**Figure 2A, 4A**). Volcano plots reveal a genome-wide decrease in the transcriptional response to *Pseudomonas* (**Figure 4B**), and finally, the *Pseudomonas* responsive genes in *set-2* animals were largely included in the control response **(Figure 4C**). Analysis of the genes two-fold downregulated in control animals shows that most of the genes were reduced at similar levels after *set-2* RNAi and were part of the same transcriptional response (**Supplemental_Figure_4A, B**). Taken together, this data suggests that *set-2* RNAi mediates part of the response to *Pseudomonas* in low SAM.

**Figure 4:**
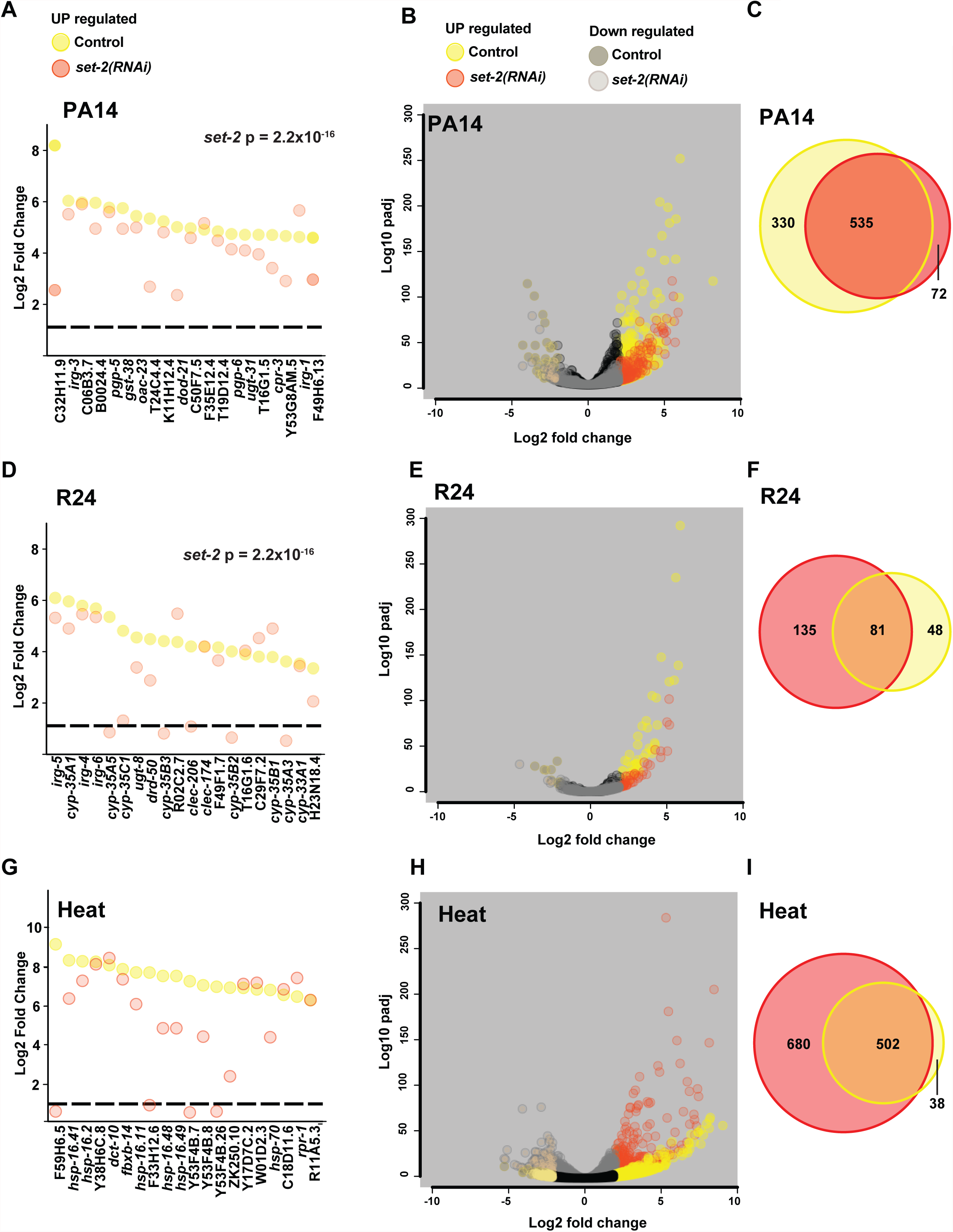
Differential transcriptional responses to a bacterial, xenotoxic and heat stress after *set-2(RNAi)* (**A**) Strip plots show that many of the top 20 genes upregulated in response to *Pseudomonas* are reduced after *set-2* RNAi. KS calculations were used to determine significance. (**B**) Volcano plots of RNA sequencing data show that fewer genes are upregulated after *set-2(RNAi)* than in controls exposed to *Pseudomonas*. (**C**) Venn diagrams show that the majority of genes upregulated after *set-2(RNAi)* in response to *Pseudomonas* were also upregulated in controls. (**D**) Strip-plot shows that many of the top 20 genes induced by R24 in control animals are reduced after *set-2* RNAi. (**E**) Volcano plots show that the transcriptional response to R24 is reduced in *set-2(RNAi)* animals compared to controls. (**F**) Venn diagram shows that *set-2(RNAi)* animals induce many genes outside the response to R24 seen in control animals. (**G**) Strip plots demonstrate that most of the top 20 genes induced in response to heat are expressed at similar levels in *set-2* RNAi animals. (**H**) Volcano plots show that more genes are significantly upregulated after *set-2(RNAi)* than in control animals in response to heat stress. (**I**) Venn diagrams show that many the majority of genes induced more than 2-fold in control animals are also upregulated after heat stress in *set-2* RNAi animals and that expression of many additional genes also increases. RNA for sequencing was isolated from control, *sams-1, set-2* and *set-16* RNAi as a set for each stress. Therefore, control genes in A-I are the same as in Figure 1 for *Pseudomonas* and Figure 2 for R24 and heat. For A-I: Control: yellow, up; brown: down; *set-2(RNAi)* orange, up; tan, down. Significant genes in RNA sequencing data (B-I) were from duplicate experiments and showed more than 2-fold changed with an FDR of <0.01.

Low SAM decreased the ability of *C. elegans* to transcriptionally respond to xenobiotic R24 (**Figure 3A-D**). Interestingly, knockdown of *set-2* mirrored *sams-1* RNAi in some respects, but not others. Like *sams-1*, heat maps show that *set-2(RNAi)* limited the number of genes upregulated by more than two-fold (**Figure 4E**). As with the response to *Pseudomonas*, *set-2* RNAi animals show reductions in several of the top 20 R24-induced genes and have diminished genome-wide expression R24 response genes (**Figure 4D, E**). However, a set of 135 genes were induced in response to R24 in *set-2* RNAi animals that were not upregulated in controls (**Figure 4F**), suggesting deregulation or expansion of the transcriptional response.

Next, we examined the response to heat stress after *set-2* RNAi and found similarities with the *sams-1* response. First, genes induced more than two fold are similar in *set-2* RNAi and controls (**Figure 3E**). Second, although most of the top 20 genes upregulated in control animals were expressed after *set-2* RNAi at near normal levels, examination of genome-wide changes showed a more extensive response (**Figure 4G, H**). Strikingly, many genes ectopic to the control response were induced after *set-2* RNAi (**Figure 4I**). As in the upregulated gene sets, the downregulated genes in control animals in response to heat also decreased after *set-2* RNAi. However, a large number of genes not downregulated in controls also decreased (**Supplemental_Figure_4E, F**). Thus, *set-2* appear to be important for full response to *Pseudomonas* or R24, but dispensable for genes induced by heat in control animals. Interestingly, knockdown of this H3K4 methyltransferase appears to deregulate or expand the stress response to both R24 and heat.

### Differential attenuation of stress-responses after knockdown of the H3K4 methyltransferases *set-16*

Like *set-2*/SET1, *set-16*/MLL is important for H3K4me3 in the *C. elegans* intestine (Ding et al. 2015). In addition, we identified a critical role for *set-16* in mediating *Pseudomonas*-responsive gene regulation in our previous studies (Ding et al. 2015). Therefore, we also compared bacterial, xenotoxic and heat stress induction in knockdown of *set-16* to *set-2* and *sams-1*. Confirming our previous qPCR analysis of selected *Pseudomonas*-responsive genes in *set-16* RNAi animals, we found that *set-16* was broadly important for expression genes upregulated by bacterial stress (**Figure 2A**). Many of the highest expressed genes in control animals during the *Pseudomonas* response were diminished after *set-16(RNAi)* **(Figure 5A**) and genome-wide comparisons showed an attenuated response compared to controls (**Figure 5B**). Finally, most of the genes upregulated by *Pseudomonas* in *set-16* RNAi animals were also upregulated in controls (**Figure 5C**). A proportion of the genes downregulated by *Pseudomonas* in control animals were also downregulated in *set-16* RNAi animals (**Supplemental_Figure_5A, B**). Thus, *sams-1*, *set-2*, and *set-16* all appear to have critical roles in regulating genes in response to bacterial stress in *C. elegans*. Responses to R24 in *set-16(RNAi)* animals largely mirrored *sams-1* knockdown but were distinct from *set-2*. Both the number of expressed genes and the levels of the highest expressed genes were significantly decreased (**Figures 3A, 5D, E**). Finally, the majority of genes that increased in *set-16(RNAi)* animals also increased in controls (**Figure 5F**). The majority of the genes downregulated by R24 in control animals were not similarly downregulated after *set-16* RNAi (**Supplemental_Figure_5 C, D**) As in *sams-1* and *set-2* knockdown, the top twenty expressed genes in control animals were expressed similarly in heat shocked control and *set-16(RNAi)* animals (**Figure 5G**) and *set-16(RNAi)* animals deregulated or expanded heat-stressed induced gene expression patterns compared to controls (**Figure 3E, Figure 5H**). Like *sams-1(RNAi)* or *set2-(RNAi)*, *set-16* animals upregulated and downregulated genes whose expression was not part of the response in control animals (**Figure 5I; Supplemental_Figure_5 E, F**). Taken together, our results suggest that low SAM, decreased *set-2*/SET1 and *set-16*/MLL activity all compromise bacterial stress-induced gene expression. Xenotoxic stress induced by R24 appeared to have a stronger requirement for *sams-1* or *set-16*, with many ectopic genes upregulated in the *set-2* response to R24. Finally, genes activated by heat shock appeared largely unaffected by low SAM, decreased *set-2*/SET1 or *set-16*/MLL activity, suggesting that neither SAM or these HMTs are essential for their expression. However, each displayed a significant number of ectopic genes inductions in both up- and down-regulated gene sets. This suggests that complex regulatory interactions may lie downstream of SAM or the H3K4 tri-methylases during the heat shock response.

**Figure 5:**
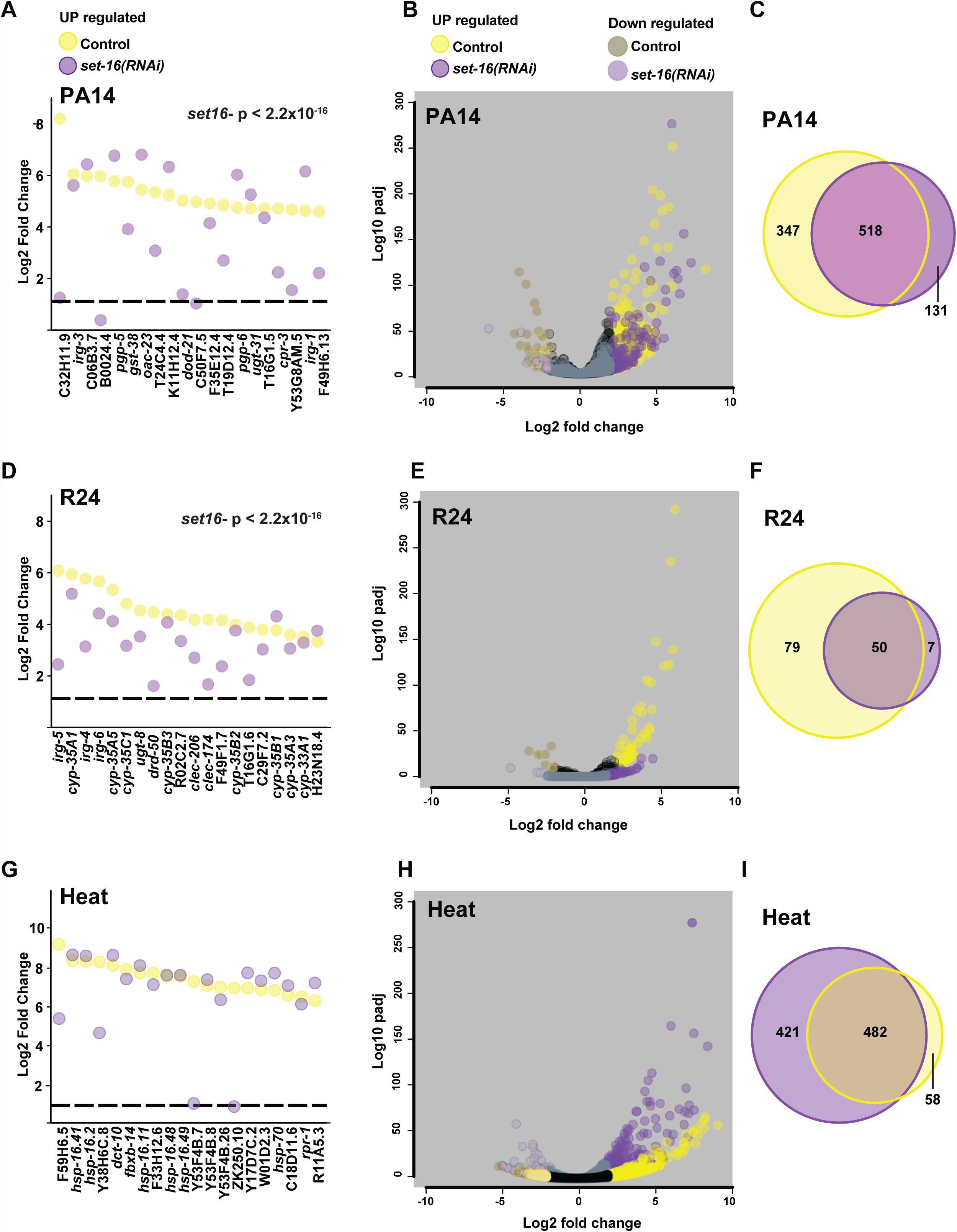
*set-16* is important for gene expression after bacterial and xenotoxic stress genes. (**A**) Strip plots show that many of the top 20 genes upregulated in response to *Pseudomonas* are reduced after *set-16* RNAi. KS calculations were used to determine significance. (**B**) Volcano plots of RNA sequencing data show that fewer genes are upregulated after *set-16(RNAi)* than in controls exposed to *Pseudomonas*. (**C**) Venn diagrams show that the majority of genes upregulated after *set-16(RNAi)* in response to *Pseudomonas* were also upregulated in controls. (**D**) Strip-plot shows that the majority of the top 20 genes induced by R24 in control animals are reduced after *set-16* RNAi. (**E**) Volcano plots show that the transcriptional response to R24 is greatly reduced after *set-16* RNAi compared to controls. (**F**) Venn diagram shows that genes induced by more than two-fold *set-16(RNAi)* animals are also induced in controls. (**G**) Strip plots demonstrate that most of the top 20 genes induced in response to heat are expressed at similar levels in *set-16* RNAi animals. (**H**) Volcano plots show that more genes are significantly upregulated after *set-16(RNAi)* than in control animals in response to heat stress. (**I**) Venn diagrams show that many the majority of genes induced more than 2-fold in control animals are also upregulated after heat stress in *set-16* RNAi animals, and that expression of many additional genes also increases. RNA for sequencing was isolated from control, *sams-1, set-2* and *set-16* RNAi as a set for each stress. Therefore, control genes in A-I are the same as in Figure 1 for *Pseudomonas* and Figure 2 for R24 and heat. For A-I: Control: yellow, up; brown: down; *set-16(RNAi)* purple, up; lavender, down. Significant genes in RNA sequencing data (BI) were from duplicate experiments and showed more than 2-fold changed with an FDR of <0.01.

### H3K4 methyltransferases *set-2* and *set-16* have differential requirements for survival during bacterial, xenotoxic or heat stress

We found that *sams-1*, *set-2*, and *set-16* were all required for the transcriptional response to bacterial stress, but differentially affected the responses to the xenotoxic agent R24. Moreover, although *sams-1*, *set-2*, and *set-16* were not required for heat shock gene expression, a varied, but a significant number of ectopic genes increased or decreased expression when knockdown animals were subjected to heat shock. Next, we sought to determine how low SAM or H3K4 HMT knockdown affected survival during each stress response. Previously, we found that *sams-1(lof)* animals had poor survival on *Pseudomonas*, which was matched by attenuated expression of bacterial-stress responsive genes, and impairment H3K4me3 acquisition at bacterial-stress responsive genes after infection (Ding et al. 2015). To determine if *set-2* or *set-16(RNAi)* animals shared this susceptibility to bacterial stress, we challenged control and knockdown animals with *Pseudomonas* and determined survival rates. Concordant with the whole genome RNA sequencing data (**Figures 4A-C; 5A-C**), we found that both *set-2* and *set-16(RNAi)* animals had significantly reduced survival on *Pseudomonas* (**Figure 6A, Supplemental_Table_4**). Our whole-genome expression analysis showed that many of the genes upregulated in control animals were part of the well-described transcriptional response to *Pseudomonas* (Troemel et al. 2006). However, we also noted other gene sets that could have important survival functions. Because GO term analysis only identifies approximately one third of annotated *C. elegans* genes (not shown), we built an annotation tool, WORM-CAT to categorize *C. elegans* genes and determine gene enrichment scores through Fisher’s exact test. WORM-CAT allows assignment of broad physiological or molecular categories (i.e., stress response), and then subsequently identifies specific sub-categories (pathogen, heavy metal, etc.) (See **Supplemental_Table_5**). If genes do not have a clear physiological function, or are pleiotropic, molecular functions were used. We validated this tool by comparison with GO analysis of our previously published microarray data from *sams-1* and *sbp-1(RNAi)* (Ding et al. 2015) (**Supplemental_Table_6**). The most significant categories, such as stress response pathogen in *sbp-1(RNAi)* and *sams-1(RNAi)* upregulated genes or fatty acid metabolic genes in *sbp-1(RNAi)* down or *sams-1(RNAi)* upregulated genes were identified in both tools. While transcriptional regulation was identified by GO ontogeny for *sams-1(RNAi)* upregulated genes, our tool showed a breakdown showing an enrichment for nuclear hormone receptors, providing additional specificity. This tool was also able to show enrichment for regulation of 1CC genes in *sbp-1* downregulated genes, which we had previously noted (Walker et al. 2011), but were not identified by GO ontogeny. Thus, this tool increases the depth and specificity of gene function in comparison to GO term enrichment.

**Figure 6:**
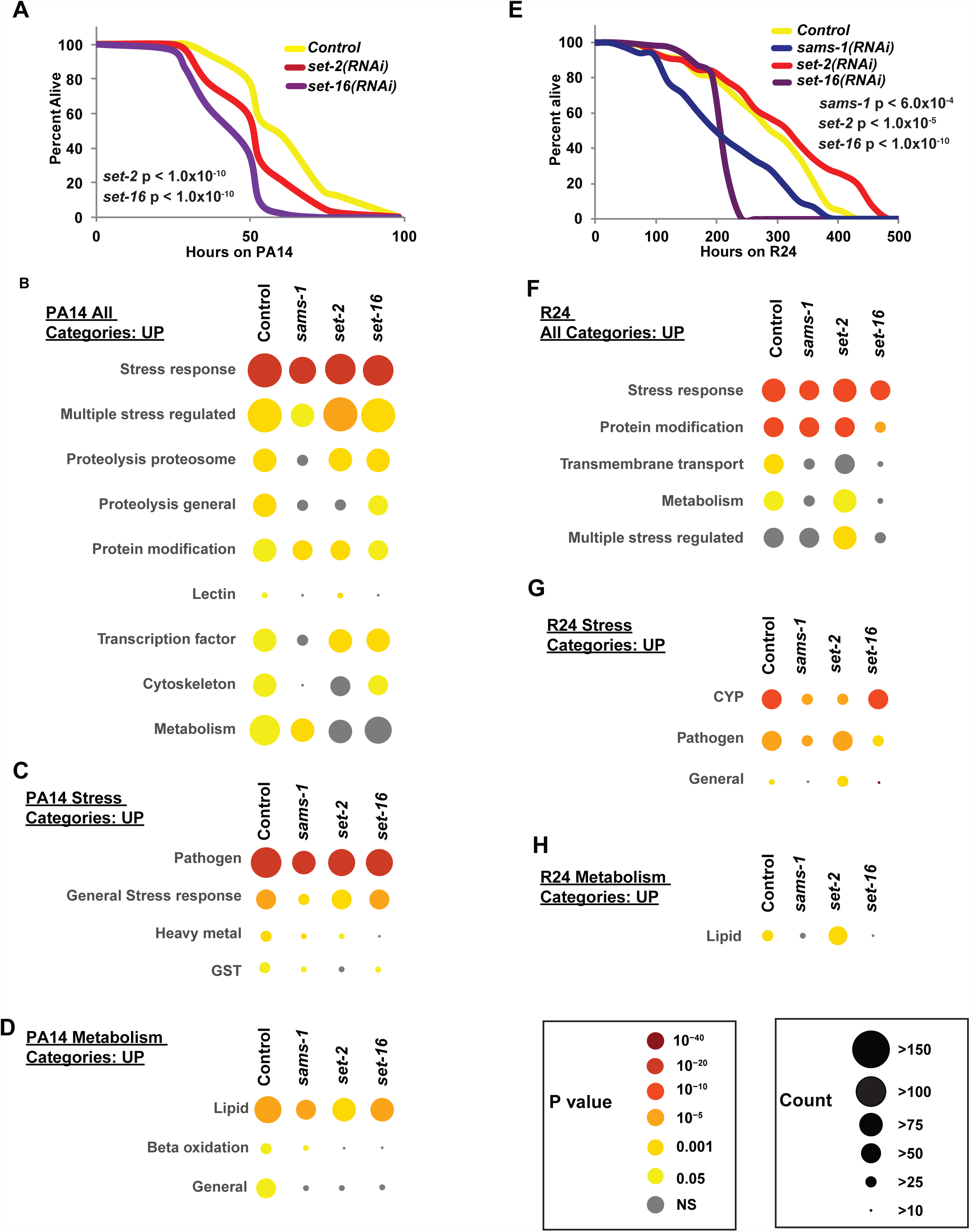
Stress-responsive and metabolic gene regulation are linked to survival after bacterial or xenotoxic stress in low SAM or H3K4 MT knockdown. (**A**) Representative Kaplan-Meier survival plot of *set-2* or *set-16* adults exposed to *Pseudomonas* shows increased sensitivity to bacterial stress (data and additional statistics available in **Supplemental_Table 4**). Statistical significance is shown by Logrank test. (**B**) Bubble charts show broad category enrichment of upregulated genes determined by WORM-CAT in control, *sams-1, set-2* or *set-16* animals in genes changed more than 2-fold (FDR <0.01) after *Pseudomonas* exposure. Specific category enrichment of stress (**C**) or metabolic (**D**) categories in *sams-1, set-2* or *set-16* animals in genes changed more than 2-fold (FDR <0.01) after *Pseudomonas* exposure. (**E**) Representative Kaplan-Meier survival plot of *sams-1*, *set-2* or *set-16* RNAi adults exposed to R24 shows differential sensitivity to xenotoxic stress (data and additional statistics available in **Supplemental_Table 8**). Statistical significance is shown by Logrank test. (**F**) Bubble charts show broad category enrichment of upregulated genes determined by WORM-CAT in control, *sams-1, set-2* or *set-16* animals in genes changed more than 2-fold (FDR <0.01) after R24 exposure. Specific category enrichment of stress (**G**) or metabolic (**H**) categories in *sams-1, set-2* or *set-16* animals in genes changed more than 2-fold (FDR <0.01) after R24 exposure.

We used WORM-CAT to determine the major categories of genes that were changed in control, *sams-1*, *set-2* or *set-16* RNAi animals, then determined which categories matched survival patterns. As expected, stress-responsive genes were fewer in number after *sams-1*, *set-2* or *set-16* knockdown (**Figure 6B**) and the stress-responsive categories were strongly enriched for pathogen responses (**Figure 6C, Supplemental_Table_7**). Enrichment scores are still highly significant, as fewer genes are upregulated in these RNAi animals. Surprisingly, metabolic genes were a significant fraction of genes upregulated in control animals and less enriched after *sams-1*, *set-2* or *set-16* RNAi (**Figure 6B**), correlating with poor survival of these animals. A breakdown of specific categories of metabolic genes showed that lipid metabolism was a significant category in control animals and that there were fewer genes and lower enrichment scores after *sams-1*, *set-2* or *set-16* RNAi. Interestingly, fatty acid desaturases have been linked to responses of *C. elegans* to *Pseudomonas* (Pukkila-Worley and Ausubel 2012). In addition, 1CC genes are among those downregulated by *C. elegans* during *Pseudomonas* infection (**Supplemental_Table_7**). Taken together, this suggests metabolic regulation may be an important part of these bacterial stress response.

R24 is a xenotoxic agent that activates immune and detoxification responses as well as aiding in the survival of *C. elegans* challenged with *Pseudomonas* but paradoxically limiting the survival of wild-type animals (Pukkila-Worley and Ausubel 2012; Pukkila-Worley et al. 2014; Cheesman et al. 2016). We found that transcriptional responses to R24 were distinct in *sams-1*, *set-2*, and *set-16* animals, with genes ectopic to the control xenotoxic stress response increasing in *set-2* RNAi animals (**Figure 3A-D; Figure 4D-F; Figure 5D-F**). To determine how *sams-1*, *set-2* and *set-16* knockdown affected survival, we treated animals with R24 and monitored death rates. In contrast to the bacterial stress survival assays, we found that *sams-1*, *set-2*, and *set-16* RNAi had distinct survival patterns. First, we found that concordant with the poor expression of xenotoxic agent-response genes, *sams-1* and *set-16* had poor survival rates, with *set-16* animals showing a particularly sharp decline (**Figure 6E, Supplemental_Table_8**). Knockdown of *set-2*, however, did not decrease survival. Next, we used WORM-CAT to identify gene categories that might correlate with sensitivity to the xenotoxic agent in *sams-1* and *set-16* animals, or survival in the *set-2* cohort. First, we noticed that as expected, stress-response genes were the most enriched category in control animals, with fewer genes in the sensitive *sams-1* or *set-16* animals (**Figure 6F, Supplemental_Table_9**). Breakdown of stress categories shows enrichment for cytochrome P450 genes and pathogen response genes (**Figure 6G**), as expected for R24 (Pukkila-Worley and Ausubel 2012). However, the knockdowns most sensitive to R24 (*sams-1* and *set-16*) differed slightly in their stress response profiles. After RNAi of *sams-1*, both CYP450 and pathogen response gene categories loose enrichment (**Figure 6G, Supplemental_Table_8**). However, *set-16* lost enrichment only within the pathogen category (**Figure 6G, Supplementary table 9**), suggesting pathogen responses are necessary for survival on R24. Supporting this notion, *set-2*, which survived normally, lost CYP450 enrichment but retained pathogen responses (**Figure 6G, Supplemental_Table_9**). We noted that metabolic categories were also limited in the sensitive strains. Metabolic genes, particularly in the lipid metabolism category were significantly enriched in Control and *set-2* RNAi animals after R24 treatment, but enrichment scores failed significance after *sams-1* or *set-16* RNAi (**Figure 6F, H, Supplemental_Table_9**). This suggests that as in the bacterial stress response, rewiring metabolic genes correlates with stress survival. Notably, an RNAi screen for R24-dependent regulators of *irg-4* identified multiple genes involved in fatty acid synthesis (Pukkila-Worley and Ausubel 2012). Finally, we found that genes deregulated in *set-2(RNAi)* animals were enriched for genes activated by multiple stresses (**Figure 6F, Supplemental_Table_9**). However, since these genes have no other functional classification, their importance of this phenotype is unclear.

### SAM and the H3K4 methyltransferases *set-2* and *set-16* are differently required for survival during heat stress

Transcription of heat shock genes in response to high temperature is controlled by shifting RNA pol II from a paused to the elongating form (Vihervaara et al. 2018). We compared heat-shock responsive transcription in low SAM or after knockdown of the *set-2*/SET1 or *set-16*/MLL methyltransferases to transcriptional changes occurring after bacterial or xenotoxic stress responses. Strikingly, we found that many genes ectopic to the control heat shock response were activated or repressed after *sams-1*, *set-2* or *set-16* RNAi (**Figure 3E-H; 4H-J; 5H-J**). Next, we performed survival assays to determine if these gene expression changes altered survival of these animals during stress. Unlike the bacterial stress response, *sams-1*, *set-2*, and *set-16* all had distinct survival curves. First, *sams-1* animals were markedly resistant during the first half of the assay, with the survival percentage at the assay midpoint more than twice that of control animals (**Figure 7A, Supplemental_Table_10**). The endpoint of the assay, however, was close to controls. Second, as in R24 assays, *set-2* animals survived most similar to controls, although p values showed a significant difference (**Figure 6A, 7A, Supplemental_Table_10**). Finally, *set-16* RNAi caused an extreme sensitivity to heat stress (**Figure 7A, Supplemental_Table_10**), similar to *Pseudomonas* and R24 responses.

**Figure 7:**
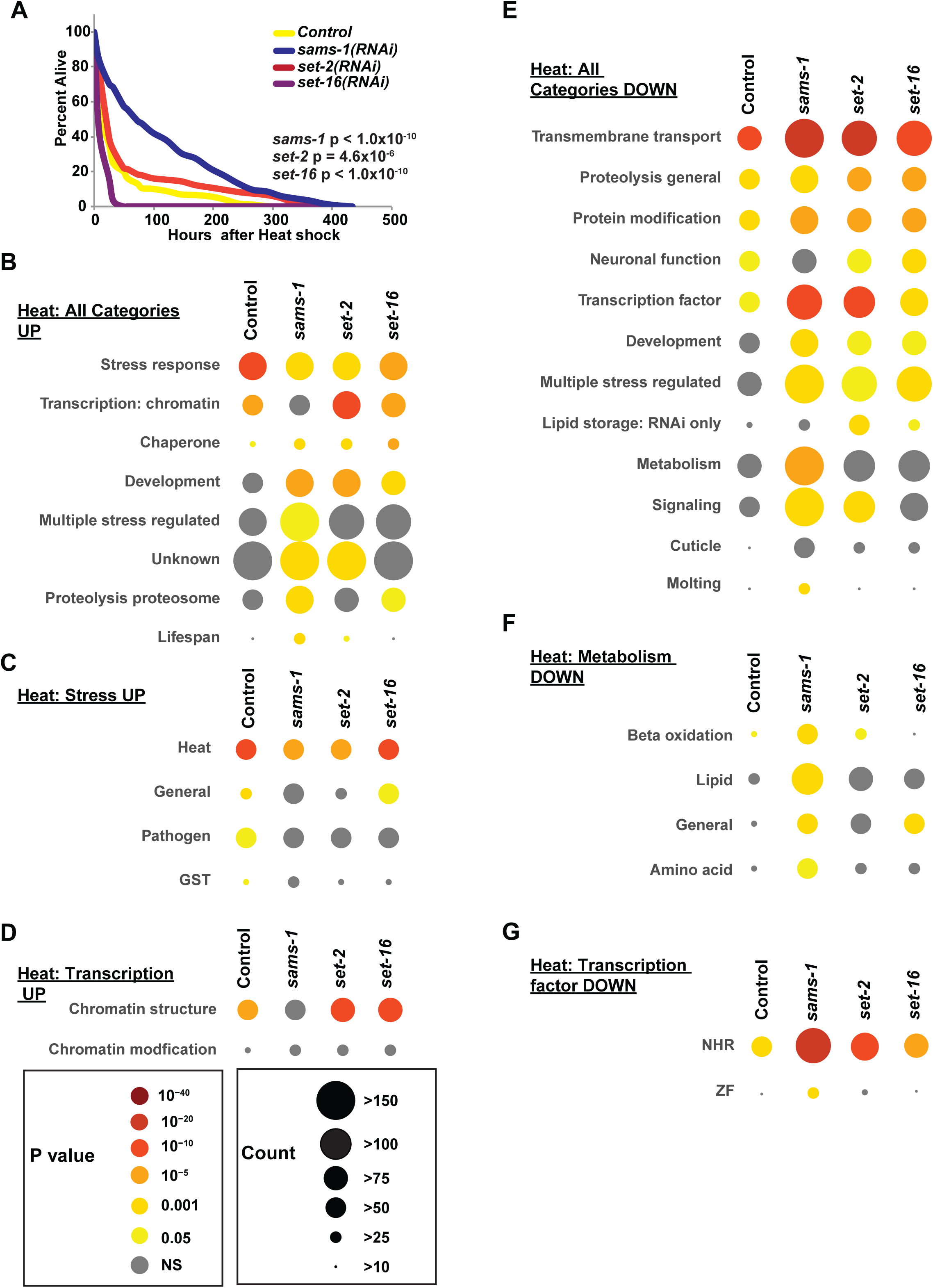
Down-regulation of metabolic gene regulation is linked to survival after heat stress in low SAM or H3K4 MT knockdown. (A) Representative Kaplan-Meier survival plot of control, *sams-1*, *set-2* or *set-16* adults exposed to heat shows increased sensitivity to heat stress (data and additional statistics available in **Supplemental_Table 10**). Statistical significance is shown by Log-rank test. for additional replicates. (B) Bubble charts show broad category enrichment of upregulated genes determined by WORM-CAT in control, *sams-1, set-2* or *set-16* animals in genes changed more than 2-fold (FDR <0.01) after heat exposure. Specific category enrichment of stress (C) or transcription (D) categories in control, *sams-1, set-2* or *set-16* animals in genes changed more than 2-fold (FDR <0.01) after heat exposure. (E) Bubble charts show broad category enrichment of downregulated genes in control, *sams-1, set-2* or *set-16* animals in genes changed more than 2-fold (FDR <0.01) after heat exposure. Specific category enrichment of metabolism (F) or transcription (G) categories in control, *sams-1, set-2* or *set-16* animals in genes changed more than 2- fold (FDR <0.01) after heat exposure.

To determine if categories of genes expressed in the *sams-1*, *set-2* or *set-16* RNAi animals correlated with the differential survival, we used WORM-CAT to survey the heat responsive genes. As expected from our initial analysis, significant numbers of stress-responsive genes were enriched in the upregulated *sams-1*, *set-2* and *set-16* RNAi cohorts (**Figure 7B, C, Supplemental_Table_11**). Genes in chromatin structure were also enriched in all but *sams-1* RNAi animals (**Figure 7B, D, Supplemental_Table_10**). However, none of these category clusters correlated with survival. Next, we examined the categories enriched in the genes downregulated during the heat shock response (**Figure 7E-G; Supplemental_Table_11**). There were several categories of enriched genes (transmembrane transport, proteolysis, and protein modification) among the *sams-1*, *set-2* or *set-16* RNAi animals (**Figure 7E, Supplemental_Table_11**). However, two categories stood out for strong enrichment in the longer-surviving *sams-1* animals, reduced enrichment in the medium surviving control and *set-2(RNAi)* animals and most attenuated in the sensitive *set-16(RNAi)* animals: metabolism and transcription factors. Strikingly, metabolism was only enriched as a downregulated category in *sams-1(RNAi)* animals during heat shock, with the majority of these genes in anabolic pathways such as lipid and amino acid metabolism (**Figure 7E, F; Supplemental_Table_11**). *C. elegans* contains a major expansion of nuclear hormone receptors, many of which are thought to regulate metabolic processes (Antebi 2015). Intriguingly, we also observed reduced NHR gene expression *sams-1(RNAi)* animals (**Figure 7G, Supplemental_Table_11**), concomitant with the loss of metabolic gene expression. Finally, we find that regulation of metabolic gene expression also correlates with survival in heat stress, as it did in our bacterial or xenotoxic stress assays, suggesting metabolic flexibility may be a common effector in stress response survival (**Supplemental_Figure_6**).

## Discussion

Metabolites that contribute to cellular regulatory functions, such as the methyl donor SAM, could be predicted to broadly affect processes such as transcription. Indeed, SAM has been proposed as a link between nutrition and regulation of the 1-carbon cycle to transgenerational epigenetic effects (Mentch and Locasale 2016). However, work from several labs across multiple systems has shown that SAM has surprisingly specific effects on histone methylation, reducing or increasing H3K4me3 as levels fall or rise (Shyh-Chang et al. 2013; Kraus et al. 2014; Ding et al. 2015; Mentch et al. 2015). Since H3K4me3 is tightly associated with start sites of actively transcribed genes, this suggests SAM may also have a critical role in acute gene regulatory events. Recently, the Locasale lab has shown that H3K4me3 peak breadth is sensitive to methionine levels in mouse liver and human cancer cells, strengthening the connections between SAM and H3K4me3 *in vivo* (Dai et al. 2018).

In this study, we have defined a role for SAM in the regulation of two stress responses, bacterial and xenotoxic stress, and found that it is necessary for induction of specific response genes as well as for survival. This link between 1-carbon metabolism and stress responses has important implications for how organisms can respond to stress when metabolically challenged. Interestingly, SAM levels had a different impact on heat shock gene expression; the heat shock response appeared normal, yet many other genes appeared to be de-repressed or ectopically activated. The release of paused RNA polymerases enables transcription of many genes in the canonical heat shock response (Vihervaara et al. 2018). Therefore SAM-dependent histone methylation may not be a primary regulator. However, Labbadia and Morimoto have recently shown that in *C. elegans*, non-cell autonomous mechanisms linked to repressive H3K27me3 limit stress responses when reproduction starts (Labbadia and Morimoto 2015). Thus, this ectopic gene activation could result from changes in repressive methylation or other indirect effects. Finally, downregulation of two classes of genes correlates with survival of *sams-1* RNAi animals under heat shock: metabolic genes in amino acid, lipid and beta-oxidation and nuclear hormone receptors. We hypothesize that downregulating these anabolic categories allows a survival advantage for *sams-1* animals as they respond to heat stress. Nuclear hormone receptors are common regulators of metabolic genes (Antebi 2015), and while direct relationships are not yet discernable, it is intriguing that both classes of these genes are downregulated in the surviving animals. Thus, low SAM may have both direct and indirect effects that influence gene expression and survival during stress responses.

SAM is utilized by histone methyltransferases such as *set-2*/SET1 or *set-16*/MLL to produce a methyl marks such as H3K4me3. Intriguingly, KTM2s are among the most sensitive histone methyltransferases to SAM levels (Mentch and Locasale 2016). SET1 is the single H3K4me3 in yeast, and thus essential for all H3K4 methylation (Shilatifard 2012). Neither *set1* or H3K4 tri-methylation are essential for viability under normal conditions (Lenstra et al. 2011). However, *set1* appears to function to limit the expression of ribosomal genes during the response to diamide (Weiner et al. 2012). The mammalian family is complex with seven H3K4 methyltransferases that differ in specificity for mono, di or trimethylation (Shilatifard 2012). However, it has been difficult to assign specific biological functions. In *C. elegans* where the KTM2 family is simpler, we have found that *set-2* or *set-16* RNAi mirrors some of the effects of low SAM, reducing the capacity to transcriptionally respond to multiple stresses. However, *set-2*/SET1 and *set-16*/MLL appeared to have distinct functional profiles during these stress responses. *set-2*/SET1 is similar to low SAM in the response to bacterial stress. Some SAM-dependent *Pseudomonas* responsive genes are also limited after *set-2* RNAi, and *set-2* animals survive poorly. Our previous analysis of the *Pseudomonas* response in *set-2* and *set-16* animals suggested that *set-2* may have a more limited role (Ding et al. 2015). The present whole genome study also bears out an essential role for *set-16* in *Pseudomonas*-responsive transcription, but notably, *set-2* animals are also survived poorly on PA14, suggesting that key genes are limited in both cases. However, *set-2* appears less critical for some genes in the detoxification response to R24. Survival is close to wild-type, and intriguingly, metabolic genes related to lipid synthesis are upregulated, distinct from the control response. Like the response to low SAM during heat shock, *set-2* RNAi did not limit expression of genes induced by heat shock in control animals while many genes were de-repressed or ectopically regulated outside the response in controls. As with R24, *set-2* RNAi animals survived similarly to controls during heat stress. Therefore changes in gene expression did not impact these biological functions. *set-2* also has intriguing functions during lifespan regulation in *C. elegans*. Greer et al. showed important transgenerational effects on lifespan in *set-2* mutants and another study from the Brunet lab suggested that *set-2* and another H3K4 HMT (*ash-2*) linked lipid synthesis and lifespan regulation (Han et al. 2017). They found that *ash-2* was important for non-cell autonomous germline to intestine regulation of these processes(Han et al. 2017). However, the role of *set-2* in direct regulation was less clear. Notably, in our study, although lipid biosynthetic genes were not changed in *set-2* animals at the L4/young adult time point, many of these genes did increase upon R24 treatment. Taken together, this suggests set-2 may impact regulation of lipid synthesis genes at different points during *C. elegans* lifespan or during specific stress responses.

*set-16*/MLL, on the other hand, was essential for survival in each of the stresses we tested. Transcriptional responses bacterial stress and R24 were attenuated, similar to low SAM. *sams-1* and *set-16* RNAi animals both survived poorly on *Pseudomonas* and R24. However, the *set-16* RNAi animals were particularly sensitive to R24. Interestingly, *set-16* animals were more deficient in activating pathogen response than CYP in response to R24 suggesting that pathogen response genes may be key for survival. Like *sams-1* and *set-2* RNAi, heat stress of *set-16(RNAi)* animals produced similar activation to control in the top 20 genes, in addition to ectopic activation or derepression of many other genes. However, this did not enhance survival. Taken together, this suggests that *set-16* has a distinct role in survival during diverse stress responses.

During a stress response, many genes must be coordinately regulated downstream of specific signaling pathways. For example, pathogenic stress may be sensed by activation of Toll-like receptors in mammals and *Drosophila* (El Chamy et al. 2015) or by translational attenuation in *C. elegans* (Troemel 2012). These signals are carried through stress-specific transcription factors that activate protective genes. Along with these direct regulatory pathways, the chromatin environment must be permissive. It is intriguing that a metabolic pathway producing the methyl donor SAM and the H3K4 methyltransferases *set-2* and *set-16* are critical to enable transcriptional responses to acute stress. This suggests that 1CC status could influence how cells or organisms could respond to outside insults. The Halsted lab, using a micropig model of alcoholic fatty liver disease, has found that dietary limitation of methyl donors markedly decreases the time for development of liver injury (Halsted et al. 2002; Medici and Halsted 2013), thus, we suggest that low SAM could exacerbate disease progressing by limiting the ability of a tissue to respond to additional stress. Finally, other metabolites such as Acetyl CoA and NAD+ also influence gene regulation (Walsh et al. 2018). By having multiple metabolic pathways influencing histone modification and gene regulation, cells might finely tune transcription to diverse nutritional signals providing templates for specific metabolic states.

## Materials and Methods

### *C. elegans* culture, RNAi and stress applications

*C. elegans* (N2) were cultured using standard laboratory conditions on *E. coli* OP50. Adults were bleached onto RNAi plates for control (L4440), *sams-1*, *set-2* or *set-16* and allowed to develop to the L4 to young adult transition before stresses were applied. RNAi efficacy was confirmed by qRT-PCR for each sample (not shown). For bacterial stress RNA preparations, nematodes were placed on *E. coli* or *Pseudomonas* plates for 6 hours. For xenotoxic stress applications animals were placed on DMSO or 70 uM R24 plates for 6 hours. For heat stress applications, animals were raised at 15°C from hatching then at the L4/young adult transition replicate plates were placed at 15°C or 37°C for 1 hour. After each stress, animals were washed off the plates with S-basal, then pellets frozen at - 80°C. RNA was prepared as in Ding, et al. 2015 (Ding et al. 2015). For survival assays, animals remained on plates until all nematodes were dead. Dead animals were identified by gentle prodding and removed each day. Kaplan-Meir curves were generated with Oasis (Yang et al. 2011).

### RNA sequencing and data analysis

RNA for deep sequencing was purified by Qiagen RNAeasy. Duplicate samples were sent for library construction and sequencing at BGI (China). Raw sequencing reads were processed using an in-house RNA-Seq data processing software Dolphin at University of Massachusetts Medical School. The raw read pairs first were aligned to *C. elegans* reference genome with ws245 annotation. The RSEM method was used to quantify the expression levels of genes (Li & Dewey, 2011, PMID: 21816040).

### Computational methods

Graphing for volcano, scatter and strip plots, Venn diagrams and bubble charts was done in R. The ontogeny category tool (WORM-CAT) consists of three parts. First, over 16,000 *C. elegans* genes were annotated; first by physiological role, then by molecular function. Categories contain up to three levels, for example, Proteolysis Proteosome: E3: F-box could appear as Proteolysis Proteosome in the broad Category 1 or as Proteolysis Proteosome: E3 or Proteolysis Proteosome: E3: F-box in the more specific categories 2 and 3. Genes with broad physiological functions (e.g., *ama-1*, RNA polymerase II large subunit) were retained in molecular function categories. Phenotype data from alleles or RNAi were used to annotate physiological role if corroborated in two or more different assays. In addition, genes with no other function whose expression was changed by at least two of these stresses (Methylmercury, tunicamycin, rotenone, cadmium, ethanol, D-glucose) were placed in the category: Stress response: regulated by multiple stresses. Annotations were applied to genes regulated in each condition, then statistical significance of category enrichment determined by Fisher’s exact test (example R script in supplemental methods) with a p-value of < 0.05 used to determine significance.

### Data access: all data will be deposited in GEO servers before publication

## Acknowledgments

We would like to thank Stefan Taubert and Jessica Feldman for critical reading of the manuscript and Thomas Fazzio and Yvonne Edwards for discussion of bioinformatics. We also thank Marian Walhout, Craig Peterson and Oliver Rando for helpful discussions. This work was supported by NIH grants R01 AG053355 (NIA) and R01 DK084352 (NIDDK).

## Author Contributions

W.D and D.Y. performed experimental work and data analysis. D.H. contributed to bioinformatics. R.P.W. provided reagents, experimental planning, data analysis and manuscript editing. A.K.W. designed and performed experiments, data analysis and bioinformatics and wrote the manuscript.

## List of supplemental information

**Supplemental Figure 1.** Gene expression changes after *sams-1* RNAi are largely PC dependent.

**Supplemental Figure 2.** Few significant changes in gene expression occur after *set-2* or *set-16* RNAi in basal conditions.

**Supplemental Figure 3.** Role of *sams-1* in genes downregulated during bacterial, xenotoxic or heat stress.

**Supplemental Figure 4.** Role of *set-2* in genes downregulated during bacterial, xenotoxic or heat stress.

**Supplemental Figure 5.** Role of *set-16* in genes downregulated during bacterial, xenotoxic or heat stress.

**Supplemental table 1:** Table showing *Pseudomonas* regulated genes (tab1: control up, tab2: *sams-1* up; tab3: *set-2* up; tab4: *set-16* up; tab5: control down; tab6: *sams-1* down; tab5: *set-2* down; tab6: *set-16* down). Genes appearing in Tromel, et al. (Troemel et al. 2006) are designated.

**Supplemental table 2:** Table showing R24 regulated genes (tab1: control up, tab2: *sams-1* up; tab3: *set-2* up; tab4: *set-16* up; tab5: control down; tab6: *sams-1* down; tab5: *set-2* down; tab6: *set-16* down).

**Supplemental table 3:** Table showing Heat shock regulated genes (tab1: control up, tab2: *sams-1* up; tab3: *set-2* up; tab4: *set-16* up; tab5: control down; tab6: *sams-1* down; tab5: *set-2* down; tab6: *set-16* down).

**Supplemental table 4:** Table showing representative PA14 survival data. Statistics generated by OASIS (https://sbi.postech.ac.kr/oasis/)(Yang et al. 2011).

**Supplemental table 5:** Table of all annotated genes with Category listing. Annotation was generated by assigning physiological function, then molecular function defined using homology, GO ontogeny and phenotype.

**Supplemental table 6:** Tables comparing WORM-CAT (tabs: *sbp-1* UP, *sbp-1* DOWN, *sams-1* UP, *sams-1* DOWN) to GO terms (tabs: GO *sbp-1* UP, GO *sbp-1* DOWN, GO *sams-1* UP, GO *sbp-1* DOWN) for previously published data from microarrays from genes changed 2 fold with a p value of < 0.05 from *sams-1(RNAi)* or *sbp-1(RNAi)* animals (Ding et al. 2015). GO term categories that are also represented in our tool are highlighted in yellow.

**Supplemental table 7:** Tables containing PA14 regulated gene category statistics for Control, *sams-1, set-2* and *set-16* RNAi (tabs show for all genes, broad, stress, metabolism for up regulated and down regulated genes). P values generated by Fisher’s exact test, significance defined as lower than 0.05. Total number of genes in categories listed in parenthesis (Column A). Pink color shows categories with a significant change in any of the RNAi animals. NS is not significant.

**Supplemental table 8:** Table showing representative R24 survival data. Statistics generated by OASIS (https://sbi.postech.ac.kr/oasis/) (Yang et al. 2011).

**Supplemental table 9:** Tables containing R24 regulated gene category statistics for *Control, sams-1, set-2* and *set-16* RNAi (tabs show for all genes, broad, stress, metabolism for up regulated and down regulated genes). P values generated by Fisher’s exact test, significance defined as lower than 0.05. Total number of genes in categories listed in parenthesis (Column A). Pink color shows categories with a significant change in any of the RNAi animals. NS is not significant.

**Supplemental Table 10:** Table showing representative heat survival data. Statistics generated by OASIS (https://sbi.postech.ac.kr/oasis/) (Yang et al. 2011).

**Supplemental Table 11.** Tables containing heat regulated gene category statistics for *Control, sams-1, set-2* and *set-16* RNAi (tabs show for all genes, broad, stress, metabolism for up regulated and down regulated genes). P values generated by Fisher’s exact test, significance defined as lower than 0.05. Total number of genes in categories listed in parenthesis (Column A). Pink color shows categories with a significant change in any of the RNAi animals. NS is not significant.

